# Folate depletion alters mouse trophoblast stem cell regulation *in vitro*

**DOI:** 10.1101/2023.09.29.558846

**Authors:** Joanna Rakoczy, Erica D. Watson

## Abstract

Maternal folate deficiency increases risk of congenital malformations, yet its effect on placenta development is unclear. Here, we investigated how folate-depleted culture medium affects the developmental potential of mouse trophoblast stem cells (TSCs). When cultured in stem cell conditions, TSC viability was unaffected by folate depletion, but ectopic differentiation of several trophoblast cell subtypes occurred. When cultured in conditions that promote differentiation, folate-depleted TSCs were driven towards a syncytiotrophoblast cell fate potentially at the expense of other lineages. Additionally, trophoblast giant cell nuclei were small implicating folate in the regulation of endoreduplication. Therefore, dietary folate intake likely promotes placenta development.

## 1. Introduction

Folate (folic acid) supplementation during pregnancy protects against fetal growth restriction (FGR) [1], though the protective mechanism is not well understood. Placental dysfunction causing FGR is likely initiated by poor development of the trophoblast lineage [2], cells that form a major part of the mature placenta. Others have linked misexpression of folate transporters and metabolic enzymes with pregnancy complications that implicate the placenta [3-7] and associated high folate concentrations with increased placental cell division and trophoblast invasion [8]. One-carbon metabolism, which includes the folate cycle, is necessary for thymidine and amino acid synthesis and the transmission of one-carbon methyl groups [9] required for epigenetic regulation of gene expression. Consequently, maternal folate intake supports processes fundamental to development. Yet, investigation into the specific effects of folate deficiency on trophoblast development is needed.

Trophoblast stem cells (TSCs), derived from trophectoderm of the blastocyst, can be maintained *in vitro* or directed to differentiate into all trophoblast cells of the mature placenta [10] **(Fig. 1A)**. By studying TSCs, researchers gain insight into the early stages of placental development and the underlying molecular mechanisms by manipulating culture conditions. Here, we analysed the morphological and molecular responses of mouse TSCs to folate-depleted (FD) culture medium with respect to stem cell maintenance and differentiation capacity.

**Fig. 1.**
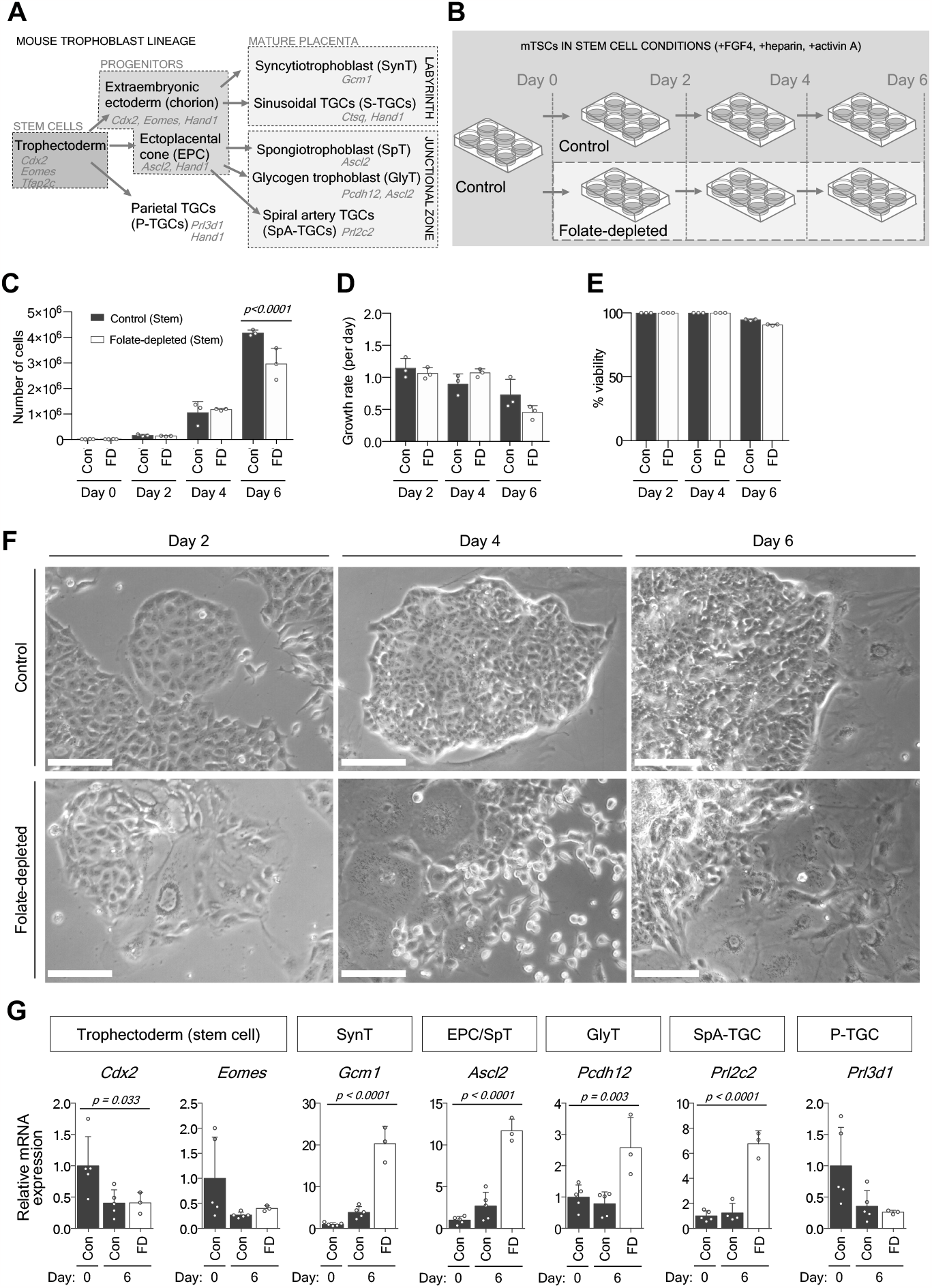
Folate depletion promotes ectopic differentiation of mouse TSCs in stem cell conditions. **(A)** Mouse trophoblast cell lineage in vivo. Gene markers are in italics. **(B)** Experimental set-up: mouse trophoblast stem cells (TSCs) were cultured for 6 days in medium promoting stem cell maintenance (contains FGF4, heparin and activin A) with normal folate (control, con) or folate-depleted (FD) levels. **(C)** Phase contrast micrographs of TSCs cultured in control and FD medium at Day 2, 4, and 6 with growth factors. **(D-F)** Characteristics of TSCs in control (dark grey bars) and FD (white bars) medium at Day 0, 2, 4, and/or 6 including **(D)** cell number, **(E)** cell growth rate per day, and **(F)** viability analysis. Two-tailed Mann-Whitney test comparing experimental groups within a specific day. *p* values indicated on graph. **(G)** RT-qPCR analysis of trophoblast marker gene expression in TSCs cultured in control (dark grey bars, Day 0 and 6) or FD (white bars, Day 6 only) medium with growth factors. Data presented as fold change (mean ± sd) relative to control mTSCs at Day 0, normalised to 1. Ordinary one-way ANOVA with Sidak’s multiple comparison test. *p* values indicated on graph.

## 2. Methods

### 2.1 Cell culture

Mouse eGFP-TSCs [10] (passage 30, from M. Hemberger, University of Calgary) were plated on polystyrene 6-well plates at 25,000 cells for 6-day cultures and 100,000 cells for Day 0 analyses. TSCs were maintained as previously described [10] including in RPMI 1640 medium (containing1.0 mg/L folic acid; ThermoFisher Scientific) with 20% fetal bovine serum (contains background folate concentration of 1.5 nmol/L [11]), 25 ng/mL fibroblast growth factor 4 (FGF4), 1 µg/mL heparin, and 5 ng/mL activin A [12]. For the FD group, RPMI 1640 medium, no folic acid (ThermoFisher Scientific) replaced RPMI 1640 medium. TSCs were differentiated by withdrawing growth factors (i.e., FGF4, heparin, activin) [10, 12]. Media was replaced every two days. Experiments were performed with technical duplicates and 3–4 biological replicates. Manual cell counts were determined with a hemocytometer. A cell growth rate calculator was used [13]. Viability was determined using a standard trypan blue exclusion assay.

### 2.2 Reverse transcription-quantitative PCR (RT-qPCR)

Total RNA was prepared using an Invitrogen RNAqueous-Micro Total RNA Isolation Kit (Thermo Fisher Scientific) for Day 0 and 2 analyses or AllPrep DNA/RNA Mini Kit (Qiagen) for Day 4 and 6 analyses. Reverse-transcription was performed with a RevertAid H Minus First Strand cDNA Synthesis Kit (ThermoFisher Scientific) using 2 µg RNA per 20-µl reaction. PCR amplification was conducted using MESA Green qPCR MasterMix Plus Assay (Eurogentec Ltd.) on a DNA Engine Opticon2 thermocycler (BioRad). Transcript levels were normalised to *Hprt1* RNA and analysed using the 2-ΔΔCt method [14]. Experiments were conducted in triplicate with 4–5 biological replicates. Primer sequences: *Ascl2, Cdx2, Eomes, Gcm1, Hand1, Tfap2c* [15], *Hprt* [16], *Pcdh12* [17], *Prl3d1* [18], *Ctsq*: forward, 5’-AGCCCGATGGAGCAGGAG, reverse, 5’-CCAAGTGCACGTTTCCAGAG, *Prl2c2*: forward, 5’-AACGCACTCCGGAACGGGC, reverse, 5’-TGTCTAGGCAGCTGATCATGCCA.

### 2.3 Statistics and Software

Statistical analyses were performed using GraphPad Prism software (version 7). RT-qPCR data were analysed by ordinary one-way ANOVA with Sidak’s multiple comparison testing. Nuclear area was measured using ImageJ (64-bit) software (version 1.48) and analysed using unpaired, two-tailed *t* tests.

## 3. Results and Discussion

To explore the effects of low folate concentration on trophoblast stem cell maintenance, mouse TSCs were cultured in control or FD media with growth factors for six days **(Fig. 1B)**. While the number of TSCs in FD conditions decreased by Day 6 compared to controls **(Fig. 1C)**, growth rate and viability were statistically normal **(Fig. 1D,E)**. However, the morphology of FD TSCs indicated that stem cell behaviour and maintenance properties were affected by low folate levels. For instance, FD TSCs displayed properties of disrupted cell-cell adhesion [19] since they appeared disorganised and rounded instead of forming tightly-adherent colonies as in controls (**Fig. 1F**). Additionally, ectopic differentiation occurred in FD medium because there were more cells with extensive cytoplasm and large nuclei **(Fig. 1F)**. We corroborated this result by RT-qPCR analysis of trophoblast marker gene expression in TSCs after exposure to FD medium for six days. Whilst stem cell markers were maintained at expected levels, FD medium caused an up-regulation of genes characteristic of terminally-differentiated trophoblast cell subtypes **(Fig. 1A,G)**. Therefore, TSCs could self-renew and maintain stem cell identity in FD conditions, but this capability lessened over time in association with ectopic differentiation. We hypothesise that folate depletion disrupts DNA [20] and/or RNA [21] methylation maintenance at trophoblast stem cell regulator genes, thus altering pre- and post-transcriptional regulation.

Next, we determined whether FD medium altered TSC differentiation potential. Trophoblast marker gene expression was assessed in TSCs exposed to control or FD media without growth factors **(Fig. 2A)**. Typical stem cell and progenitor marker gene expression was observed **(Fig. 2B)** confirming that stem cell potential was lost in control and FD conditions. However, FD medium (without growth factors) appeared to shift TSC differentiation towards a syncytiotrophoblast cell fate as determined by high *Gcm1* transcript levels relative to controls **(Fig. 2B)**. Simultaneously, *Ctsq* and *Prl2c2* genes failed to up-regulate by Day 6 and *Prl3d1* mRNA was prematurely up-regulated at Day 4 **(Fig. 2B)** suggesting that differentiation of several trophoblast giant cell (TGC) subtypes was disrupted by folate depletion **(Fig. 1A)**. In support, morphological analysis of FD cells revealed fewer TGCs **(Fig. 2C)**. TGCs that were present contained significantly smaller nuclear area **(Fig. 2D)**. We propose two hypotheses (not necessarily mutually exclusive): 1) folate depletion drives TSC differentiation towards a syncytiotrophoblast cell fate at the expense of TGC subtypes. This might result from altered DNA methylation causing misexpression of genes responsible for lineage decisions [22]; 2) TGCs form yet DNA endoreduplication [23] is repressed by low folate (and consequently, low thymidine [24]) levels resulting in altered amplification and transcription of TGC-specific genes [25].

**Fig. 2.**
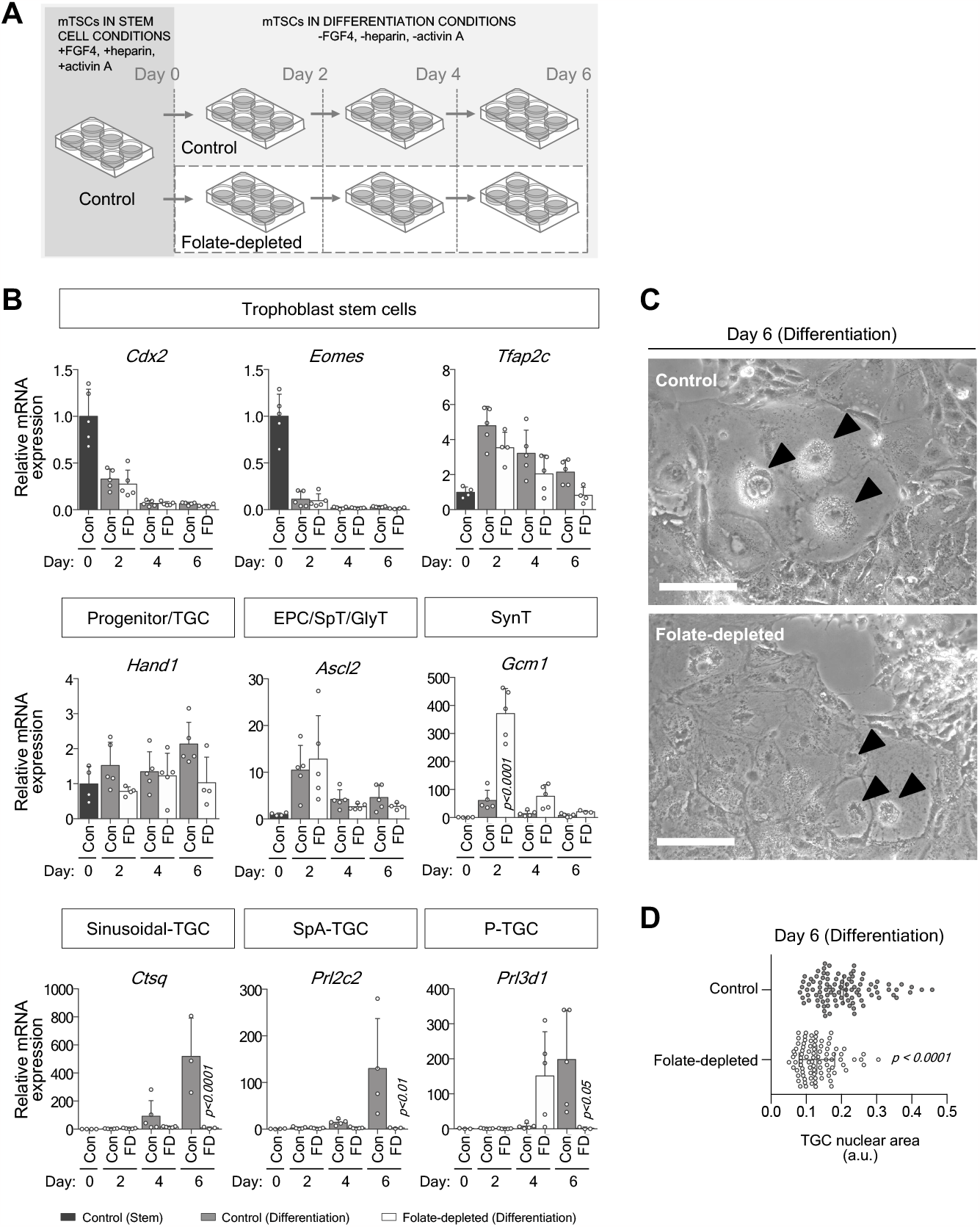
Folate depletion alters differentiation potential of mouse TSCs when growth factors removed. **(A)** Experimental set-up: mouse trophoblast stem cells (TSCs) were cultured overnight in stem cell conditions (Day 0) and then growth factors (FGF4, heparin and activin A) were withdrawn for 6 days to promote differentiation in medium containing normal folate (control, con) or folate-depleted (FD) levels. **(B)** RT-qPCR analysis of trophoblast marker gene expression in TSCs cultured in normal folate (light grey bars) or FD (white bars) levels at Day 2, 4, or 6 days after growth factor withdrawal. Data presented as fold change (mean ± sd) relative to control TSCs in stem cell conditions (Day 0, dark grey bars), normalised to 1. Ordinary one-way ANOVA with Sidak’s multiple comparison test. *p* values indicated on graph. **(C)** Phase contrast micrographs of TSCs in control and FD medium after 6 days without growth factors. Arrowhead, trophoblast giant cell (TGC) nucleus. **(D)** Quantification of mean (± sd) TGC nuclear area of differentiated TSC culture in control and FD medium. Cells were identified by the following characteristics: mononuclear, extensive cytoplasm, large nucleus. Technical duplicates, *n* = 88–97 cells. Unpaired, two-tailed *t* test. *p* value indicated on graph.

Overall, defects in trophoblast stem cell maintenance and differentiation caused by low folate concentrations have implications for placenta development and function, and by extension fetal growth.

## Abbreviations

FD: folate-depleted;
FGR: fetal growth restriction;
TSC: trophoblast stem cell;
TGC: trophoblast giant cell.

## Author Contributions

E.D.W. and J.R. conceived the project, designed the experiments, and analysed the data. J.R. performed the experiments and collected the data. E.D.W. interpreted the results and wrote the manuscript. All authors read and edited the manuscript.

## Acknowledgements

J.R. was supported by a Royal Society Newton International Fellowship (RG79568). This work was funded by the Lister Institute for Preventative Medicine to E.D.W (RG78687).

